# NXP800 suppresses HSF1 through coordinated stress signaling and GCN2-dependent translational repression in glioblastoma

**DOI:** 10.64898/2026.11.17.764637

**Authors:** Haddie De Hart, Julie Heldman, Matthew O’Malley, Andy Lian, Kirk Staschke, Richard L. Carpenter

**Author notes:** Corresponding Author Contact Information Richard Carpenter, PhD Address: 308 Biology Building, 1001 E. 3^rd^ St., Bloomington, IN 47405. **Financial Support:** National Cancer Institute (R01CA285967, RLC) and The William J. Wright Scholarship Fund at IU Simon Comprehensive Cancer Center (HD).

## Abstract

Under states of proteotoxic stress, Heat Shock Factor 1 (HSF1) serves as the primary transcription factor responsible for restoring protein homeostasis. General Control Nonderepressible 2 (GCN2) similarly alleviates cellular stress. Activated by a shortage in amino acids, GCN2 triggers the phosphorylation of eIF2α to downregulate global protein synthesis while selectively translating survival-promoting proteins like Activating Transcription Factor 4 (ATF4). Cancer cells often exploit these pathways to mitigate damage from rapid proliferation and metabolic demands. In Glioblastoma (GBM), HSF1 promotes tumor growth and apoptosis evasion, while GCN2 maintains cancer cell homeostasis and survival. NXP800 has recently emerged as both an HSF1 inhibitor and GCN2 agonist with potent antitumor activity, yet the mechanism by which these pathways integrate to modulate GBM fate remains unclear. Herein, we show NXP800 induces an initial, GCN2-independent downregulation of HSF1 transcriptional activity, followed by a delayed, GCN2-p-eIF2α-dependent decrease in HSF1 translation. This temporal regulation is accompanied by altered phosphorylation of the HSF1-activating kinase AKT1. Furthermore, we show NXP800 directly engages the HSF1 protein and that HSF1 suppression is important for the NXP800 anti-proliferative effects. Analysis of cell line and patient data further establishes a negative correlation between GCN2 and HSF1 signaling across multiple cancer types, including GBM. Collectively, our data provide a molecular framework for the dual inhibitory action of NXP800 and offers a comprehensive understanding of how this compound impairs cancer progression.

## Introduction

Maintenance of protein homeostasis is critical for cellular viability and function. Under proteotoxic stress, such as heat, oxidative damage, or protein misfolding, heat shock factor 1 (HSF1) acts as the primary transcription factor orchestrating the heat shock response (HSR) through transcriptional regulation of heat shock proteins (HSPs). In the absence of stress, HSF1 is sequestered in the cytoplasm by these same heat shock proteins. Upon activation, these chaperones release HSF1 to facilitate the refolding of denatured proteins, prevent aggregation, and target misfolded proteins for degradation. This allows HSF1 to move into the nucleus, trimerize, bind to heat shock elements, and regulate gene expression of those targets (1, 2).

Historically, HSF1 has been difficult to therapeutically target due to its lack of a stable ligand-binding pocket and dynamic protein structure (2, 3). NXP800 (CCT361814) is a compound that was originally discovered when searching for inhibitors to HSF1 (4, 5). An exciting breakthrough occurred when NXP800 entered into a Phase 1 clinical trial (NCT05226507) in patients with advanced cancers, as this was the first-in-human trial of a compound that can target HSF1. In recent years, however, it has become increasingly clear that NXP800 can target other pathways, namely it appears to agonize the integrated stress response (ISR) pathway with heavy involvement of the kinase General Control Nonderepressible 2 (GCN2) (4–9).

In times of metabolic stress, the GCN2 arm of the ISR is triggered by a depletion in amino acids. Activated by an accumulation in uncharged tRNAs, GCN2 phosphorylates the alpha subunit of Eukaryotic Initiation Factor 2 (eIF2). As a result, global translation is downregulated while favoring translation of a select group of mRNAs, such as Activating Transcription Factor 4 (ATF4). ATF4 coordinates a transcriptional program targeting genes such as DNA damage-inducible transcript 3 (*DDIT3* aka CHOP), Asparagine Synthetase (*ASNS*), and Tribbles Pseudokinase 3 (*TRIB3*) which collectively reprogram metabolism to enhance amino acid synthesis, nutrient transport, and autophagy (2, 10).

Studies focused on the intersection of HSF1 and GCN2 stress response pathways found ATF4 and its canonical targets are upregulated in response to heat shock (11), whereas inhibition of HSP70 induced an autophagy response via the GCN2-p-eIF2α pathway (12). Conversely, amino acid deprivation, the canonical activator of GCN2, results in decreased HSF1 protein levels as well as impaired HSF1 DNA-binding activity (13). Despite these observations, the precise molecular crosstalk between these two stress response pathways remains unclear.

Cancer cells frequently exploit the HSF1 and GCN2 pathways to sustain rapid proliferation and survive metabolic stress (1, 14, 15). In Glioblastoma (GBM), the deadliest form of brain cancer in adults (16), HSF1 activity is elevated in 74% of patient samples compared to normal tissue, where increased expression of HSF1 targets correlates with poor clinical outcomes (17, 18).

Furthermore, HSF1 facilitates GBM growth, migration, and apoptosis evasion, identifying it as a promising therapeutic target (17–21). GCN2 is overexpressed in approximately 90% of human cancers and is implicated in maintaining tumor homeostasis (15, 22–24). While GCN2 inhibition reduces *VEGFA* expression and angiogenesis, pharmacological activation of GCN2 has been shown to inhibit growth and induce cell death (25, 26). However, the role of GCN2 in GBM remains poorly defined.

Despite the clinical progression of NXP800 in other malignancies, its efficacy and underlying molecular mechanism in GBM remain uncharacterized. Here, we demonstrate that NXP800 activates GCN2 signaling while concurrently suppressing the HSF1 pathway in GBM. More specifically, NXP800-mediated HSF1 inhibition occurs through two distinct mechanisms. First, through an acute GCN2-independent suppression of HSF1 transcriptional activity, potentially mediated by altered protein-protein interactions or direct ligand engagement. Second, via a sustained, GCN2-dependent depletion of HSF1 protein synthesis. Finally, our work shows HSF1 suppression is important to the anti-proliferative effects of NXP800 and that the relationship between the GCN2 and HSF1 pathways extends beyond NXP800. Collectively, our data provide a molecular framework for the dual inhibitory action of NXP800 and identifies a therapeutic opportunity for the treatment of GBM.

## Results

### NXP800 suppresses HSF1 transcriptional activity prior to the depletion of HSF1 protein

NXP800 is a promising therapeutic candidate recognized for its activation of the GCN2 pathway, potent inhibition of the HSF1 pathway, and sustained anti-tumor activity both *in vitro* and *in vivo* (4, 6, 8, 27–29). While NXP800 has been investigated across several cancer types, its effects in GBM and the molecular basis of HSF1 pathway suppression remain poorly understood. To address this gap in knowledge, NXP800 was evaluated for whether it modulates GCN2 and HSF1 activity in GBM cells. Results indicated that NXP800 significantly increased the expression of the ATF4 transcriptional targets DDIT3 (CHOP) and TRIB3, confirming GCN2 pathway activation, in both U118 and T98G GBM cells (Fig. 1A-B). Conversely, NXP800 significantly reduced HSPA1A and HSPE1 mRNA levels (Fig. 1A-B), consistent with reports that NXP800 inhibits HSF1 (4, 6, 8, 28, 29). Notably, NXP800 treatment did not significantly alter HSF1 mRNA expression, suggesting that the observed pathway inhibition occurs post-transcriptionally. Under conditions of proteotoxic stress, HSF1 binds to heat shock elements (HSEs) within the promoters of target genes to drive transcriptional activation (1). Although previous studies have relied on changes in HSF1 target gene expression as a surrogate measure of pathway activity, the specific step at which NXP800 disrupts HSF1 signaling has not been investigated. Therefore, we directly assessed HSF1 transcriptional activity using a luciferase reporter containing four tandem HSE repeats. NXP800 significantly reduced HSF1-dependent reporter activity as early as 4 hours following treatment, with suppression maintained through 24 hours in all three GBM cell lines examined (Fig. 1C-E). Given the reduction in HSF1 activity without significant changes to HSF1 mRNA expression, we next investigated whether the loss of pathway signaling results from a decrease in total HSF1 protein abundance. Western blot analysis revealed NXP800 simultaneously increased ATF4 protein levels while decreasing HSF1 protein levels in both a dose-and time-dependent manner (Fig. 1F-I). Notably, HSF1 transcriptional activity declined as early as 4 hours following treatment, preceding detectable decreases in total protein levels. While a slight decrease in HSF1 protein was observed at 8 hours, consistent loss across GBM cell lines occurred at 16 hours (Fig. 1F-I). These findings suggest NXP800 suppresses HSF1 signaling through at least two distinct events, including an early inhibition of HSF1 transcriptional activity followed by a delayed depletion of HSF1 protein. This temporal separation also suggests that the initial suppression of HSF1 activity likely occurs independent of HSF1 protein loss.

**Figure 1:**
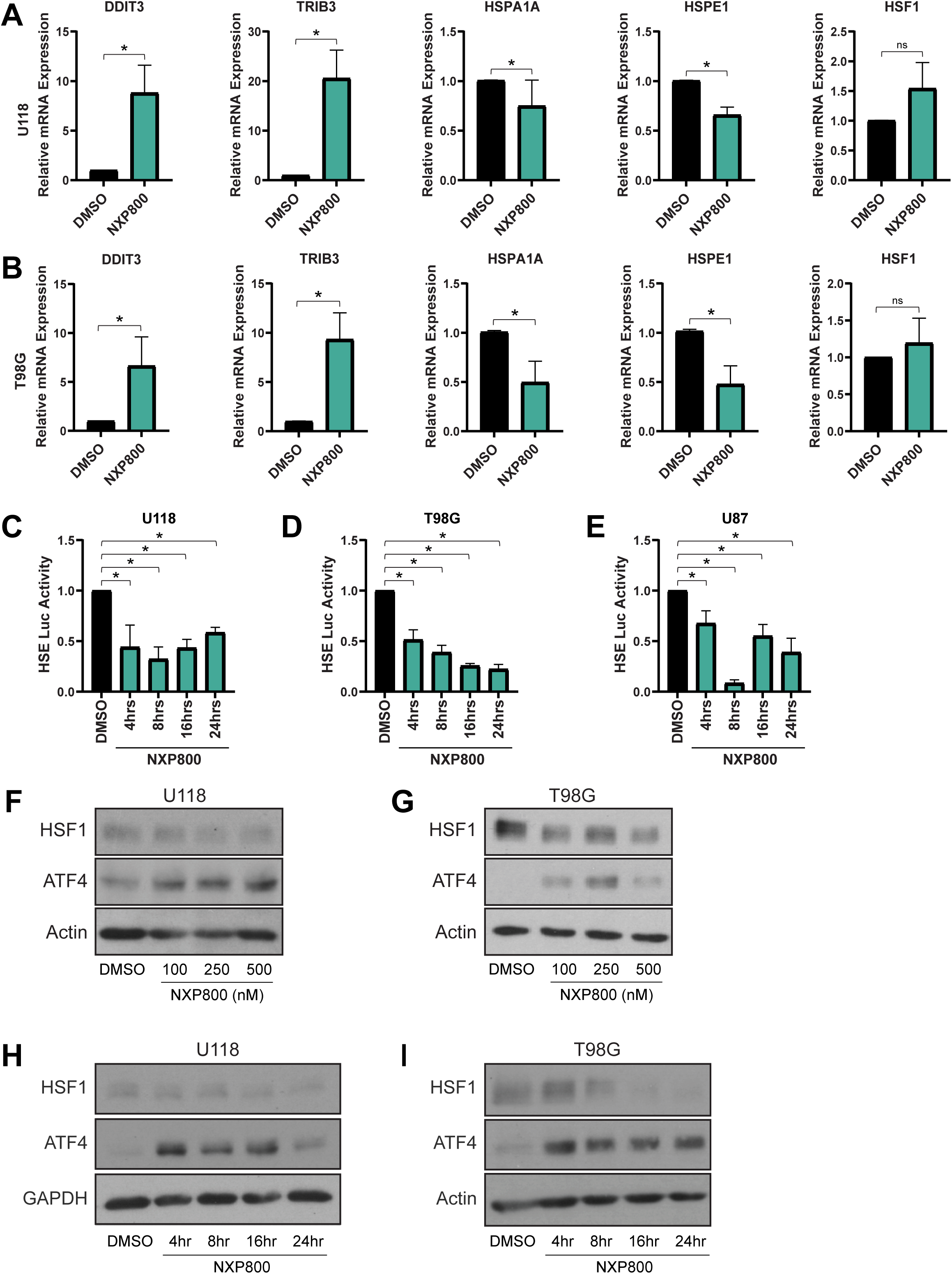
NXP800 decreases HSF1 pathway activation and HSF1 protein levels. A-B) U118 (A) and T98G (B) cells were treated with NXP800 (250nM) for 24 hours. Total RNA was collected and subjected to RT-qPCR using indicated primers. C-E) U118 (C), T98G (D), and U87MG (E) cells were transfected with pGL4-HSE and pRL-TK luciferase reporters for 24 hours. Cells were then treated with NXP800 for the indicated times. Cells were lysed and subjected to dual luciferase reporter assay as indicated in the methods. F-H) U118 (F, H) and T98G (G, I) cells were treated with NXP800 in a dose response (F-G) or time course (H-I) manner at the indicated time points and doses. The time course used 250nM NXP800. Cells were lysed and total protein subjected to immunoblotting for indicated antibodies.

### NXP800 alters activity of HSF1 activation through AKT

NXP800 suppresses HSF1 transcriptional activity prior to depletion of HSF1 protein (Fig. 1), suggesting that the early inhibition of HSF1 activity is mediated by a mechanism distinct from those regulating HSF1 protein abundance. Since NXP800 has been reported to activate the GCN2 pathway (8) and NXP800 was found to increase expression of genes downstream of ATF4 signaling (Fig. 1A), we tested whether HSF1 activity was dependent on GCN2. GBM cells were treated with NXP800 in the presence or absence of the GCN2 inhibitor, GCN2iB. NXP800 significantly reduced HSF1 activity but inhibition of GCN2 did not alter this effect, suggesting the inhibition of HSF1 transcriptional activity is independent of GCN2 (Figure 2A-C). To identify NXP800-regulated signaling pathways that could potentially alter HSF1 activity, U118 and T98G cells were exposed to vehicle or NXP800 (250nM) for 2 or 4 hours. Total RNA was then subjected to RNA sequencing and bioinformatic analysis. Principal component analysis (PCA) confirmed samples clustered with their respective treatments (Suppl. Fig 1A-B). Differentially expressed genes (DEGs) for 2 hours (Fig. 2D-E) and 4 hours (Suppl Fig 1C-D) are illustrated using volcano plots that indicate important genes that are up- and down-regulated by NXP800 as well as heat maps including all DEGs (Suppl. Fig 1E). In both cell lines, there was significant downregulation of several heat shock protein genes, suggesting HSF1 activity was decreased. To more specifically assess this, a published gene signature for HSF1 activity (30) was used to assess the effect of NXP800 and showed a significant decrease at both 2 and 4 hours in both cell lines (Fig. 2F-G). This result suggests NXP800 is effective at reducing HSF1 transcriptional activity as early as 2 hours. To identify potential pathways affected by NXP800, gene ontology was performed on both up- and downregulated genes. Interestingly, analysis of the downregulated genes indicated a decrease in AKT-mTOR signaling (Fig. 2H-I). AKT1 has previously been shown to phosphorylate S326 on HSF1 (31, 32), a key residue regulating transcriptional activity of HSF1 (32, 33). Using a published gene signature for AKT1 activity, AKT1 activity was found to be decreased by NXP800 (34) (Fig. 2J-K). Furthermore, the AKT1 signature was also decreased in prostate cancer (8) (Suppl Fig 1F). and osteosarcoma (28) (Suppl. Fig 1G-H) cells treated with NXP800, indicating this effect on AKT activity is likely a generalized effect of NXP800. Immunoblotting further showed concurrent suppression of active AKT (S473) and reduced HSF1 S326 phosphorylation as early as 1 hour post-treatment (Fig. 2L). These results suggest NXP800 inhibits AKT1 activity leading to a suppression of HSF1 transcriptional activity in as little as 1-2 hours after NXP800 treatment.

**Figure 2:**
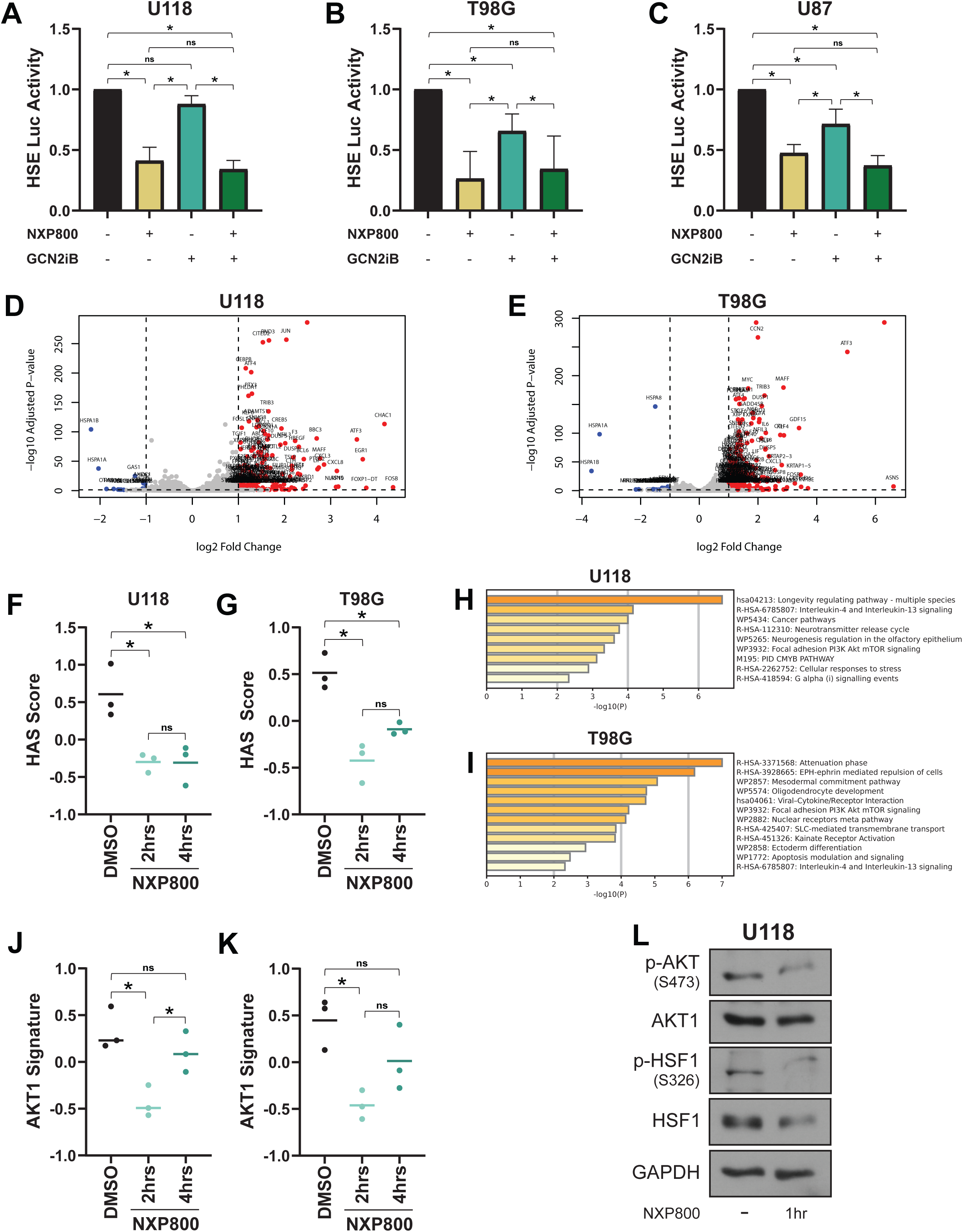
NXP800 suppresses HSF1 activity through inhibition of AKT, independent of GCN2. A-C) U118 (A), U87MG (B) and T98G (C) cells were transfected with pGL4-HSE and pRL-TK luciferase reporters for 24 hours. Cells were then treated with either DMSO, NXP800 (250nM), and/or GCN2iB (2 µM). Cells were lysed and subjected to the dual luciferase reporter assay as indicated in Materials and Methods. D-I) U118 and T98G cells were treated with NXP800 (250nM) for 2 or 4 hours and total RNA was subjected to RNA sequencing. Differentially expressed genes at 2 hours of treatment were plotted on volcano plots (D-E). Scores for the HSF1 activity signature were plotted across groups for both cells (F-G). Significantly downregulated genes by NXP800 were subjected to gene ontology analysis to identify signaling pathways altered by NXP800 (H-I). AKT1 signature scores were plotted across the groups to indicate the effect of NXP800 on AKT1 activity (J-K). L) U118 cells were treated with NXP800 (250nM) for 1 hour and total protein was subjected to immunoblotting with the indicated antibodies.

### NXP800 reduces HSF1 protein levels through GCN2-mediated translational inhibition

NXP800 has been characterized as a potent activator of the integrated stress response (ISR); however, the specific upstream kinase responsible for this effect remains a subject of ongoing investigation. While NXP800 is commonly described as a GCN2 agonist (28, 29), recent studies have proposed that NXP800 activates the ISR through PKR-like ER kinase (PERK) (8, 28).

Because subsequent experiments rely on pharmacological inhibition of GCN2, we first sought to determine the specific ISR kinase responsible for NXP800-mediated signaling. To do so, an ATF4-luciferase reporter assay was utilized in HEK293 cells. Treatment with NXP800 resulted in robust activation of the ATF4 reporter, comparable to the established GCN2 agonist halofuginone (HF) (Fig. 3A). To distinguish between the contributions of GCN2 and PERK, ATF4-Luc reporter cells were treated with NXP800 in the presence of either GCN2iB or HC- 5404, a PERK inhibitor. Results revealed potent inhibition of NXP800-induced ATF4 reporter activity by GCN2iB (IC50 = 55.1nM), whereas PERK inhibition failed to attenuate reporter activity even at high concentrations (IC50>20 μM) (Fig. 3B). These findings indicate that NXP800-mediated activation of the ISR occurs predominantly through the GCN2-ATF4 signaling axis.

**Figure 3:**
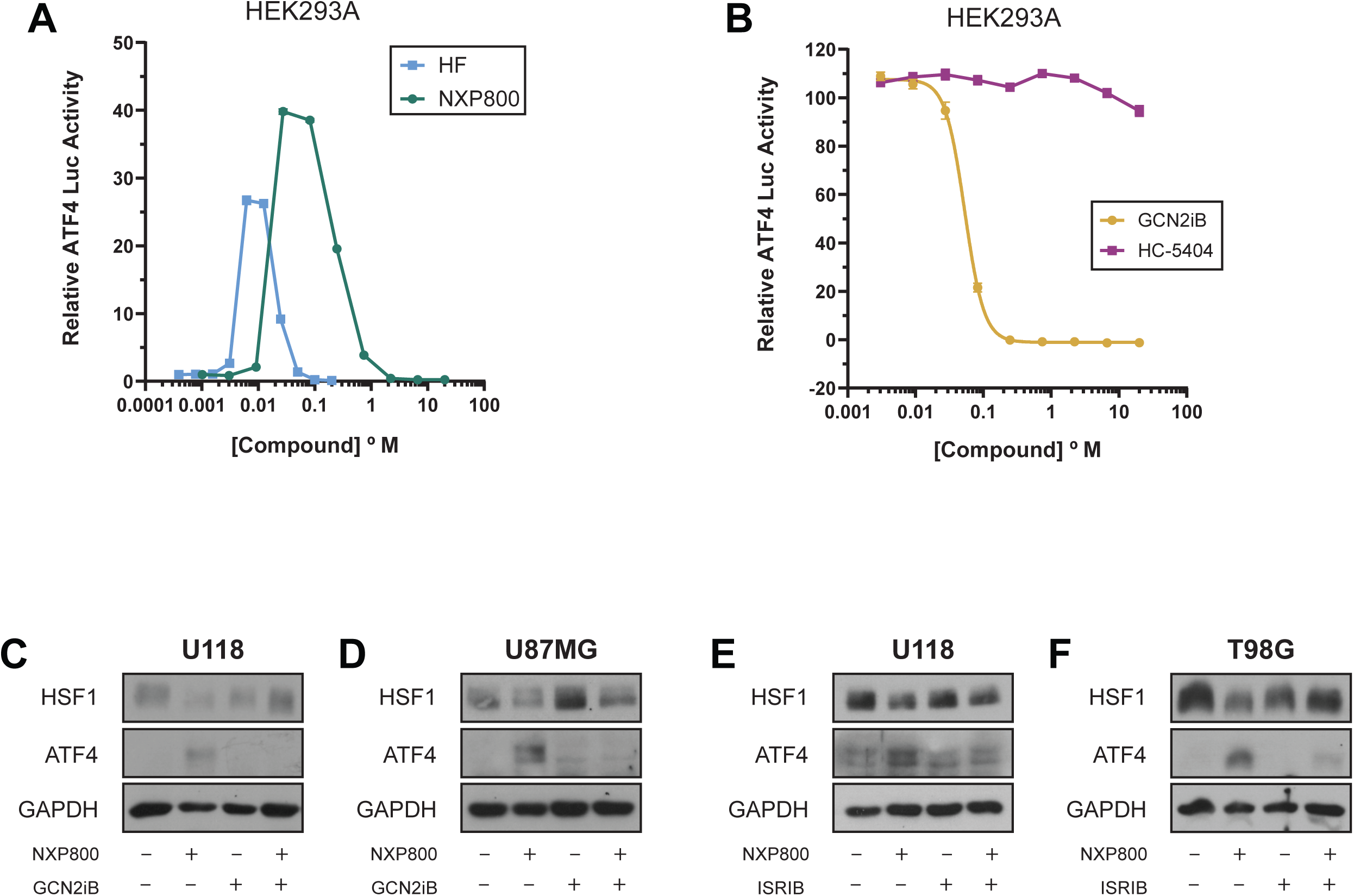
NXP800 promotes loss of HSF1 protein through GCN2-eIF2α-mediated translation suppression. A-B) HEK-293 cells stably expressing an ATF4 luciferase reporter were treated with the indicated compounds. Cells were treated with increasing doses of NXP800 or Halofuginone (HF) (A). Cells were treated with 30nM NXP800 combined with increasing doses of GCN2iB or HC-5404 (B). C-D) U118 (C) and U87MG (D) cells were treated with NXP800 (250nM) and/or GCN2iB (2 µM) for 24 hours. Total protein was subjected to immunoblotting for the indicated antibodies. E-F) U118 (E) and T98G (F) cells were treated with NXP800 (250nM) and/or ISRIB (1 µM) for 24 hours. Total protein was subjected to immunoblotting for the indicated antibodies.

Activation of GCN2 suppresses global protein synthesis through phosphorylation of eIF2α, which inhibits eIF2B activity and promotes selective translation of ATF4. The small molecule Integrated Stress Response Inhibitor (ISRIB) allosterically stabilizes eIF2B, allowing translation to proceed despite the presence of phosphorylated eIF2α (35). Having established GCN2 as the primary mediator of NXP800-induced ISR activation, we next investigated whether the delayed loss of HSF1 protein results from activation of the canonical GCN2-p-eIF2α pathway. First, we tested whether inhibition of GCN2 rescued the NXP800-induced decrease in HSF1 protein levels. Inhibition of GCN2 with GCN2iB resulted in a partial rescue of HSF1 protein levels suggesting NXP800 activation of GCN2 contributes to the downregulation of HSF1 (Fig. 3C-D). Next, cells were treated with NXP800 alone or in combination with ISRIB. Consistent with inhibition of ISR signaling, ISRIB blocked NXP800-induced ATF4 accumulation and rescued HSF1 protein levels (Fig. 3E-F). Together, these findings indicate that NXP800-mediated depletion of HSF1 protein occurs through canonical GCN2-p-eIF2α ISR signaling and support a model in which activation of translational repression contributes to the loss of HSF1 protein.

### NXP800 engages HSF1 protein

While prior work suggests GCN2 is a target of NXP800 (28), we show NXP800 suppresses HSF1 activity independent of GCN2 activity. This suggests NXP800 may engage other proteins beyond just GCN2. To determine whether NXP800 physically engages HSF1, we performed a thermal shift assay with NXP800 and the direct HSF1 inhibitor SISU-102 (DTHIB) (36) as a positive control. Following treatment, cell lysates were exposed to a thermal gradient followed by immunoblotting to analyze the remaining soluble fractions. Results showed that NXP800 enhanced HSF1 thermal stability relative to DMSO (Fig. 4A-B). The shift in the HSF1 melting profile suggests NXP800 may engage the HSF1 protein and warrants further study to identify the direct targets of NXP800.

**Figure 4:**
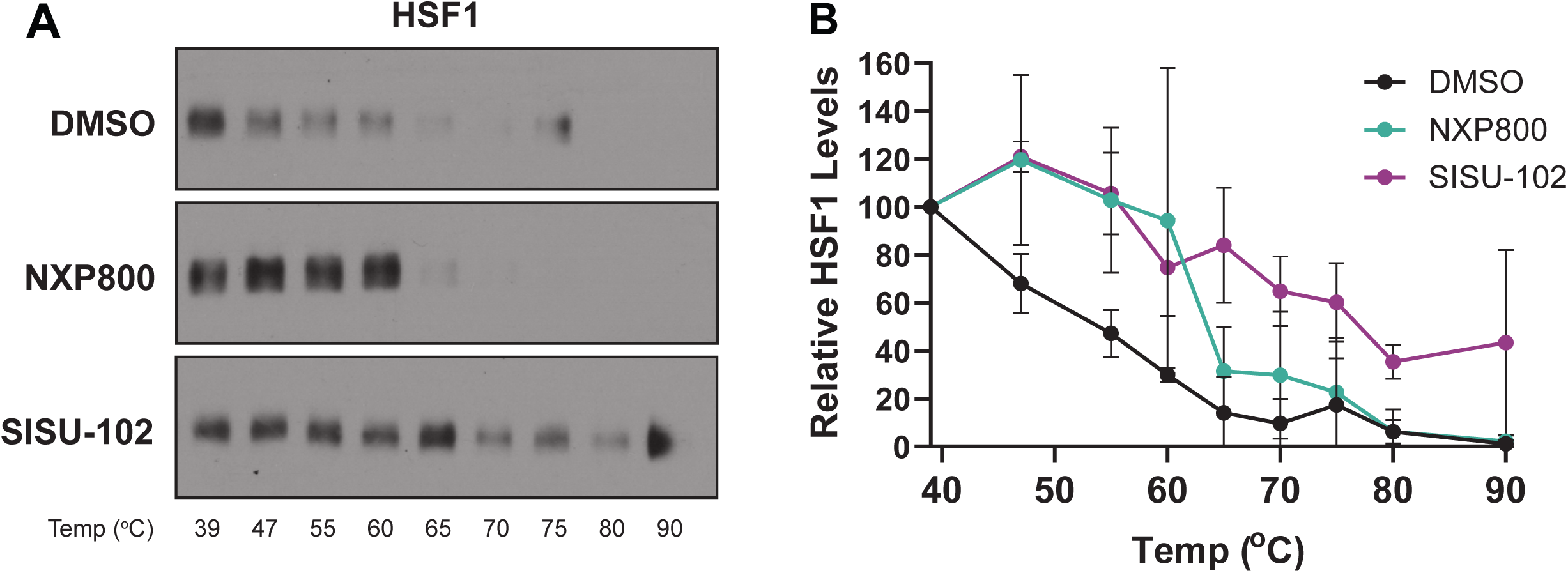
NXP800 engages the HSF1 protein. A) U118 cells were subjected to the target engagement assay as described in the methods in the presence of DMSO, NXP800 (250nM), or SISU-102 (5 µM). B) Signal was quantified across increasing temperatures.

### NXP800 reduced GBM cell growth that was partially dependent on HSF1

NXP800 has been reported to delay tumor growth, inhibit proliferation, and stunt clonogenic capacity in ovarian cancer, prostate cancer, osteosarcoma and hepatocellular carcinoma (6, 8, 28, 29). To evaluate the anti-tumor effect of NXP800 in GBM, a clonogenic assay was performed and found NXP800 significantly decreased colony forming ability in both U118 and T98G (Fig. 5A-B). Furthermore, NXP800 was observed to decrease cell viability over time with significant reductions in viability being observed after 16-48 hours of treatment (Fig. 5C-F). As HSF1 is well documented to play a role in promoting malignant growth in cancer (3, 14), it was next determined whether HSF1 was critical for the NXP800 anti-growth effect. To do so, we subjected previously established control or HSF1 knockdown breast cancer cells (37) to treatment with either DMSO or NXP800 for 24-48 hours. As expected, knockdown of HSF1 or NXP800 treatment alone reduced cell viability (Fig. 5G). Although NXP800 continued to reduce cell viability in HSF1-deficient cells, the additional inhibitory effect was no longer statistically significant (Suppl. Fig. 2), the additional inhibitory effect was no longer statistically significant (Fig. 5G). These findings indicate HSF1 contributes to the anti-proliferative activity of NXP800 but is not solely responsible for its effects, consistent with NXP800 acting through multiple signaling pathways.

**Figure 5:**
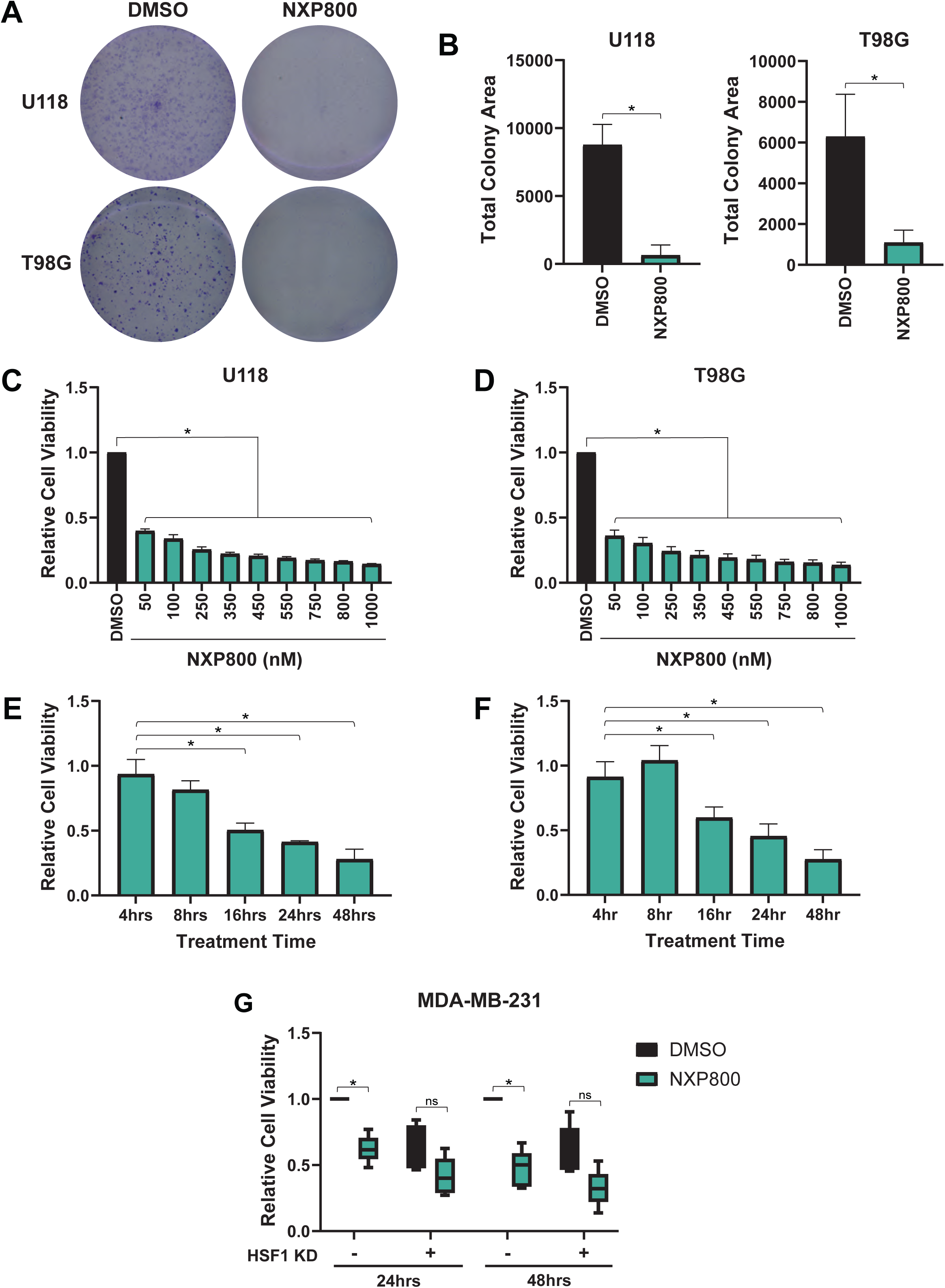
NXP800 suppresses the growth of GBM cells that is partially dependent on HSF1. A-B) U118 and T98G cells were subjected to clonogenic growth assays in the presence or absence of NXP800 (250nM) (A). Area covered by colonies was quantified using FIJI software (B). C-F) U118 (C, E) and T98G (D, F) cells were subjected to a cell viability assay in the presence of NXP800 for either (C-D) 48hrs at the indicated doses or (E-F) with 250nM of NXP800 for the indicated time points. G) MDA-MB-231 cells with control or HSF1 knockdown were subjected to cell viability assays in the presence or absence of NXP800 (250nM) for the indicated time points.

### GCN2 Activation is Negatively Correlated with HSF1 Activation

GCN2 activation is triggered by amino acid deprivation, inducing GCN2 autophosphorylation and the subsequent phosphorylation of eIF2α. This signaling axis drives the selective translation of the transcription factor ATF4, which orchestrates a transcriptional program to mitigate cellular stress (2, 10, 38, 39). Due to the post-translational regulation of this pathway, GCN2 mRNA expression is often decoupled from its enzymatic activity, making GCN2 transcript levels an unreliable method of determining GCN2 activity. To provide a more robust assessment of GCN2 activity in patient data, we utilized a computational approach to identify a gene set whose expression positively correlates with GCN2 activation.To establish this subset of GCN2- responsive genes, we analyzed publicly available RNA-seq data from fibroblast-like synoviocytes treated with the GCN2 agonist Halofuginol (HFol), a derivative of the prolyl-tRNA synthetase inhibitor Halofuginone (HF) (40). This analysis initially yielded a list of 386 significantly upregulated genes. To isolate GCN2-specific targets, we excluded genes that retained HFol-responsiveness during GCN2 depletion, resulting in a 17-gene list hereafter referred to as the GCN2 Activity Signature (GAS) (Fig. 6A-C). To further validate this signature as a representation of GCN2 activity, publicly-available data (41) was used to assess the response of the GAS gene list to GCN2iB. The GAS showed significant decreases with 6 and 24 hours of GCN2iB treatment (Fig. 6D, Suppl. Fig. 4A), indicating the GAS can detect changes with pharmacological targeting of GCN2. The pan-ErbB kinase inhibitor Neratinib was recently shown to directly interact with and activate GCN2 (25). To continue validating the GAS genes, they were assessed for their responsiveness to neratinib. Results indicated the GAS was increased in response to neratinib treatment for 72 hours but this response did not occur with GCN2 knockout (Fig. 6E, Suppl. Fig. 4B). Lastly, the GAS was assessed for response to NXP800 from a previous study (8). Treatment with NXP800 led to a dose-dependent increase in the GAS (Fig. 6F). Together, these results confirm the GAS as a measure of GCN2 activity across different activation mechanisms and in different cellular contexts.

**Figure 6:**
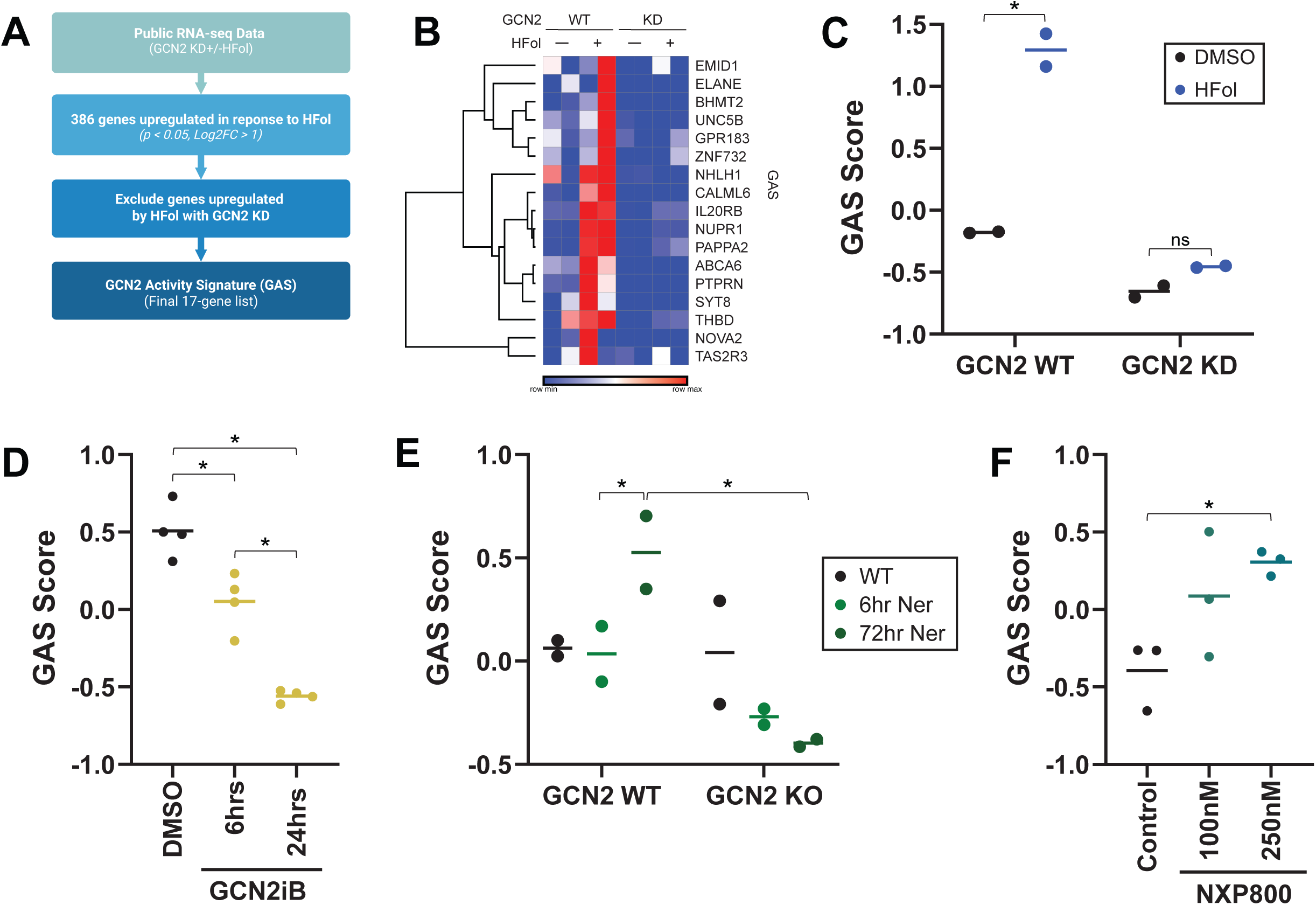
Generation of a GCN2 activity gene signature. A) Workflow to identify the GCN2 activity signature (GAS) is presented in a flow chart. B-C) Genes included in the GAS are presented via heat map (B) and using signature scores (C) from a publicly available dataset (GSE145203) that included Halofuginol (HFol) with or without GCN2 KD. D) The GAS was validated in a publicly available dataset (GSE196252) that included GCN2iB treatment at the indicated time points. E) The GAS was validated in a publicly available dataset (GSE188958) that included treatment with neratinib with or without GCN2 knockout. F) The GAS was validated in a publicly available dataset (PRJEB81176) that included treatment with NXP800 at the indicated doses.

These results suggest that agonizing the GCN2 pathway leads to a downregulation of HSF1 (Figs. 1-3), which supports previous studies showing the effects of NXP800 treatment (4, 6, 8, 28, 29). To further characterize the relationship between GCN2 and HSF1, we used the previously mentioned HAS, representing HSF1 activity, to examine changes in HSF1 activity following treatment with various GCN2 agonists. First, it was observed that the HAS was reduced with halofuginol treatment (40), which was lost with knockout of GCN2 (Fig. 7A). The HAS was also reduced by neratinib (25), which was also prevented by GCN2 knockout (Fig. 7B). Next, the HAS was found to decrease in response to NXP800 from a previous study in prostate cancer cells (8) (Fig. 7C). These results suggest agonizing the GCN2 pathway leads to an inverse effect on HSF1 activity. To assess if this relationship is maintained in GBM patients, the TCGA cohort of GBM patients was analyzed and HAS was found to be significantly negatively correlated with the GAS (Fig. 7D). This relationship was also maintained in the TCGA cohort of lower grade gliomas (LGG) (Fig. 7E), suggesting this relationship may be part of the development of low and high grade gliomas. Since NXP800 has also been shown to be effective in prostate cancer models (8), the TCGA-PRAD cohort was also assessed and this relationship was consistent in prostate cancer wherein HAS was negatively correlated with GAS (Fig. 7E).

**Figure 7:**
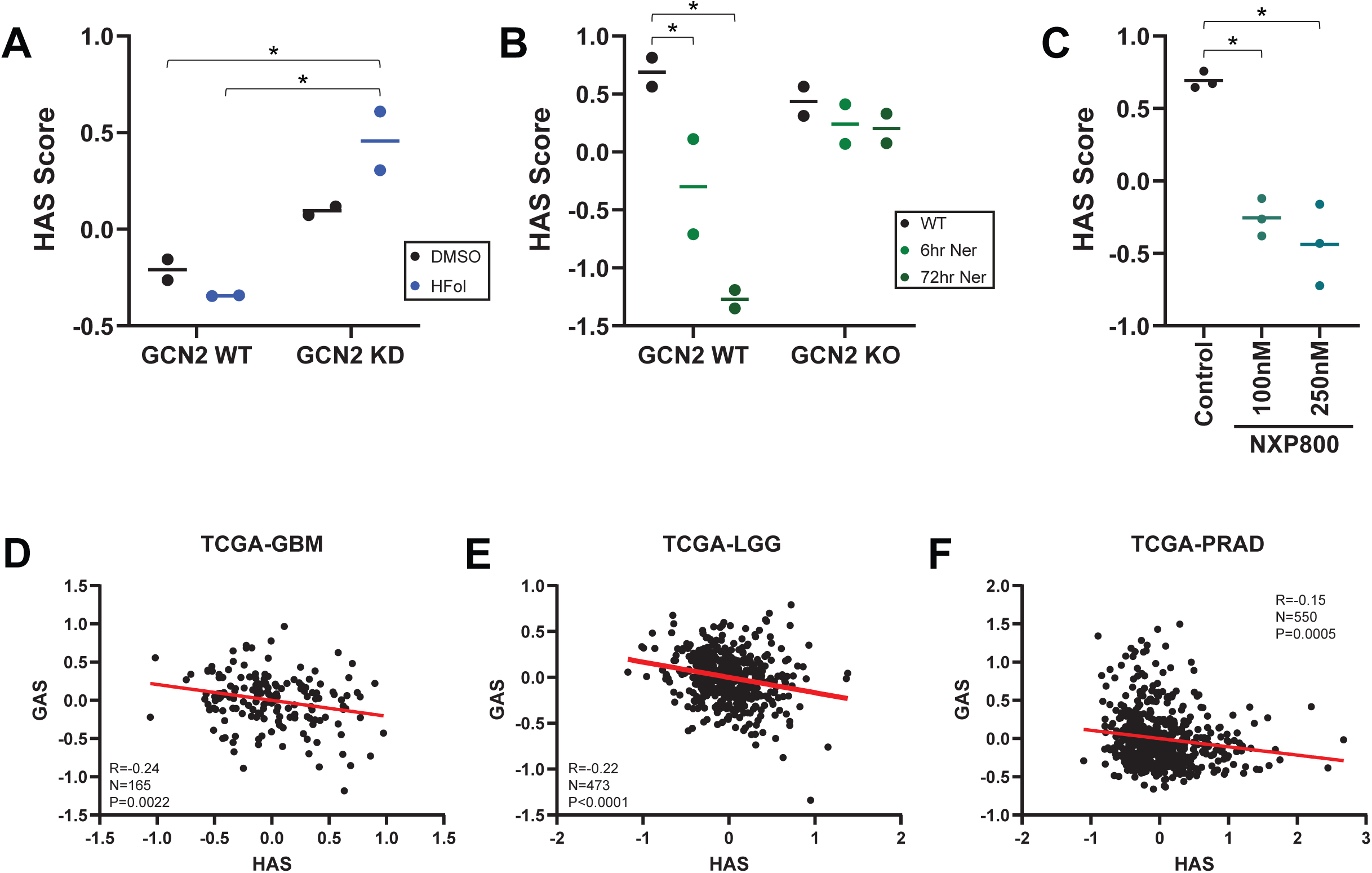
HSF1 activity is inversely related to GCN2 activity. A) HSF1 activity signature (HAS) was analyzed in a publicly available dataset (GSE145203) that included Halofuginol (HFol) with or without GCN2 KD. B) HAS was analyzed in a publicly available dataset (GSE188958) that included treatment with neratinib with or without GCN2 knockout. C) HAS was analyzed in a publicly available dataset (PRJEB81176) that included treatment with NXP800 at the indicated doses. D-F) Signature scores for HAS and GAS were correlated in TCGA cohorts for GBM (D), low grade gliomas (E), and prostate cancer (F).

These findings suggest that NXP800 influences both the GCN2 and HSF1 pathways through effects on distinct components of each pathway, as well as potential crosstalk between these pathways in the patient populations studied.

## Discussion

The established role of HSF1 in tumorigenesis and cancer progression has positioned it as an attractive therapeutic target and driven the development of small molecules capable of suppressing HSF1 signaling. In response to this need, the chemical precursor of NXP800 (CCT361814) was originally identified in a phenotypic screen for its ability to inhibit expression of the HSF1 target gene *HSPA1A* (4). Subsequent studies demonstrated that NXP800 also activates the integrated stress response through GCN2, leading to inhibition of cell proliferation and robust anti-tumor activity in multiple cancer models (8, 28, 29). These promising preclinical findings have supported the advancement of NXP800 into clinical evaluation for cholangiocarcinoma (42). Although NXP800 has therefore been widely regarded as both an HSF1 inhibitor and a GCN2 agonist, the mechanistic relationship between these activities has remained largely undefined.

Our findings support a revised model for the mechanism of action of NXP800. Interestingly, previous studies have implicated different signaling pathways in mediating the anti-tumor effects of NXP800. In prostate cancer, NXP800 has been proposed to inhibit proliferation through PERK-mediated activation of the unfolded protein response (8), whereas studies in osteosarcoma identified GCN2 as the primary mediator of its cytotoxic effects (28). Our findings provide a framework that reconciles these seemingly disparate observations by demonstrating that NXP800 coordinately perturbs multiple stress-response pathways whose relative contributions may vary depending on cellular context and the timing of pathway activation. By examining the temporal response to NXP800, we demonstrate that suppression of HSF1 signaling occurs through mechanistically distinct early and late events. Early after treatment, NXP800 suppresses HSF1 transcriptional activity independent of GCN2, with unbiased transcriptomic analysis identifying AKT signaling as one of the earliest pathways altered by NXP800 and biochemical studies confirming rapid inhibition of AKT activity. In contrast, prolonged exposure to NXP800 activates canonical GCN2-ISR signaling, resulting in depletion of HSF1 protein through translational repression. Furthermore, although HSF1 contributes to the anti-proliferative effects of NXP800, loss of HSF1 alone does not fully account for the biological response to treatment. Together, these findings support a model in which NXP800 coordinately perturbs multiple interconnected stress-response pathways that collectively suppress HSF1 signaling.

This revised mechanistic model has important implications for both therapeutic development and experimental design. From a therapeutic perspective, simultaneous modulation of multiple stress-response pathways may be the underlying reason for the potent anti-tumor activity observed for NXP800 across multiple cancer types and supports its continued clinical investigation. Conversely, therapeutic utility and mechanistic specificity are distinct properties of small molecules. Because NXP800 alters multiple signaling pathways that independently contribute to its biological effects, phenotypes observed following NXP800 treatment cannot be interpreted as selective perturbation of either HSF1 or GCN2 signaling without complementary genetic or orthogonal pharmacologic approaches. Accordingly, while NXP800 remains a promising therapeutic agent, our findings suggest it should not be considered a pathway- selective pharmacologic probe for dissecting HSF1 or GCN2 biology.

One additional finding further illustrates the complexity of NXP800 biology. We found that NXP800 altered the thermal stability of HSF1, suggesting that HSF1, or an HSF1-associated protein complex, may represent an additional molecular target of NXP800. Interestingly, in 2026, NXP800 was also suggested to interact with GCN2. These results suggest NXP800 either targets multiple proteins individually or in a complex. While thermal shift assays alone cannot distinguish between direct ligand binding and indirect stabilization through protein-protein interactions, these findings raise the possibility that NXP800 modulates HSF1 through an additional mechanism beyond kinase signaling and ISR activation. Defining the direct molecular target(s) of NXP800 will therefore be an important direction for future studies.

Beyond defining the mechanism of NXP800, our findings also provide insight into the biological relationship between the GCN2 and HSF1 stress-response pathways. Previous studies demonstrated that amino acid deprivation suppresses HSF1 activity (13), yet the mechanistic relationship between these pathways has remained largely unexplored. We show that three structurally distinct GCN2 agonists (NXP800, HFol, neratinib) each suppress HSF1 signaling while activating the GCN2 pathway. Furthermore, the inverse relationship between GCN2 and HSF1 activity was maintained across GBM and prostate cancer patient cohorts, suggesting that this interaction extends beyond pharmacologic perturbation and may represent a broader feature of tumor biology. One possible explanation for this relationship is that activation of GCN2 reduces the requirement for HSF1-mediated proteostasis by suppressing global protein synthesis, thereby shifting the cellular response from maintaining protein folding to limiting protein production during metabolic stress. Although additional studies are needed to define the molecular basis of this crosstalk, our findings support a model in which GCN2 and HSF1 function as interconnected stress-response programs rather than independent signaling pathways.

One of the principal findings of this study is the identification of AKT signaling as a candidate mediator of the early suppression of HSF1 activity by NXP800. Phosphorylation of HSF1 at S326 by AKT1 is a well-established driver of HSF1 transcriptional activity in both the heat shock response and cancer (31–33); however, whether NXP800 altered upstream regulators of HSF1 had not previously been investigated. Using early transcriptomic profiling followed by biochemical validation, we identified AKT as one of the earliest pathways altered by NXP800, with suppression of AKT signaling preceding the loss of HSF1 transcriptional activity. These findings provide a plausible explanation for the temporal separation between early inhibition of HSF1 activity and the later GCN2-dependent depletion of HSF1 protein. Although our data strongly supports a role for AKT in this early response, the mechanism by which NXP800 rapidly suppresses AKT activity remains unknown. Previous kinome profiling did not identify AKT as a direct target of the chemical precursor of NXP800 (4), suggesting that inhibition of AKT occurs indirectly through an upstream signaling event. Defining the molecular target(s) responsible for this rapid suppression of AKT signaling will therefore be an important direction for future investigation.

An important consideration for the development of HSF1-directed therapeutics is the contrasting role of HSF1 in cancer and the central nervous system. Whereas sustained HSF1 activity promotes tumor growth and survival, HSF1 is also essential for maintaining neuronal protein homeostasis and protecting against protein aggregation (3, 43). Consequently, inhibition of HSF1 in brain tumors presents a unique therapeutic tradeoff: suppressing HSF1 may limit glioblastoma progression while potentially compromising proteostasis in normal neural tissue.

Whether NXP800 crosses the blood-brain barrier remains unknown. This property would be advantageous for treating GBM but could also increase the risk of unintended effects on healthy brain tissue. Conversely, if NXP800 is developed for non-CNS malignancies, brain penetration could expose normal neurons to chronic HSF1 suppression. Future studies defining the blood- brain barrier permeability and neurological safety profile of NXP800 will therefore be important for its continued clinical development.

In summary, our study provides a revised mechanistic framework for understanding the biological activity of NXP800. Rather than functioning as a selective inhibitor of HSF1 or a simple agonist of GCN2, NXP800 coordinately modulates multiple interconnected stress- response pathways that converge on suppression of HSF1 signaling through temporally distinct mechanisms. This integrated model reconciles previously disparate observations regarding the mechanism of action of NXP800 while providing insight into the broader relationship between the HSF1 and GCN2 stress-response pathways. Collectively, our findings support the continued development of NXP800 as a promising anti-cancer therapeutic while emphasizing that its biological effects should be interpreted within the context of coordinated stress-network modulation rather than selective perturbation of a single pathway.

## Experimental Procedures Cell Culture and Reagents

Human Glioblastoma cell lines U118, U87 & T98G (ATCC) and human breast cancer cell line MDA-MB-231 (ATCC; RRID:CVCL_YJ68) were maintained in Dulbecco’s Modification of Eagle’s Medium with 4.5g/L glucose, L-glutamine & sodium pyruvate (DMEM; Corning; 10-013-CV) supplemented with 10%fetal bovine serum (FBS; Corning; 35-076-CV) and 1%Penicillin Streptomycin (PS; Corning; 30-002-CI). All cell lines were routinely maintained at 37∘C in a 5% CO2 incubator. Trypsin EDTA (1X; Corning; 25-053-CI) and DPBS (1X; Corning; 21-030-CV) were used for cell passaging and washing, respectively. Generation of shScramble or shHSF1 MDA-MB-231 are as previously described (37).

## Chemicals

NXP800 was a kind gift from Nuvectis Pharmaceuticals (New Jersey, USA). GCN2 inhibitor, GCN2iB was purchased from MedChemExpress (#HY-112654). HF (#13370), ISRIB (#41879), and SISU-102 (#37433), were purchased from Cayman Chemical. Stock solutions were prepared in 100% dimethyl sulfoxide (DMSO) and stored at -20 ◦C. NXP800 was stored at - 80◦C.

## Real-time quantitative RT-PCR (RT-qPCR)

Total RNA was extracted from cells using the Quick RNA-microprep kit (Zymo Research; ZR1051) according to the manufacturer’s instructions. cDNA was synthesized from total RNA using the High-Capacity RNA-to-cDNA kit (Applied Biosystems/Thermo Fisher Scientific; 4387410BF5). Quantitative PCR was performed on a real-time PCR system (QuantStudio 3, Applied Biosystems) using PowerUP SYBR Green (Applied Biosystems; A25743). Gene expression levels were normalized to GAPDH and quantified using the ΔΔCt method. Primers for gene targets are listed in Supplemental Table 1.

## Dual-Luciferase Assay

Cells were seeded in triplicate in a 12-well plate and then transfected 24 hours later with firefly luciferase reporter, pGL4.41[luc2P/HSE/Hygro] (Promega; E375) and renilla luciferase reporter, pRL-TK (. After 24 hours of transfection, cells were either treated with DMSO or the indicated drug. Post treatment, cells were washed in PBS then collected and measured following the Renilla Luciferase Assay System (E2810) and the Luciferase Assay Reagent ie. Firefly (E1483) protocol from Promega. Relative promoter activity was determined by normalizing the Firefly luciferase activity over the Renilla luciferase activity.

## Protein Isolation and Immunoblotting

Cells were harvested in and washed with cold Phosphate-Buffered Saline (DPBS; Corning; 21- 030-CV) and lysed in IP lysis buffer (50 mM Tris, 150 mM NaCl, 1 mM EDTA, 1% NP-40, 1% sodium deoxycholate) with HALT™ Protease and Phosphatase Inhibitor Cocktail (Pierce, Cat. #1861281). Samples were sonicated and then centrifuged to remove cellular debris, and the total protein extract was collected. Protein concentration was determined using a colormetric assay (Protein Assay Dye; Bio-rad; 5000006) according to the manufacturer’s instructions.

Proteins were subjected to Sodium Dodecyl Sulfate-Polyacrylamide Gel Electrophoresis (SDS- PAGE) on an 8 or 10% polyacrylamide gel. Samples were then transferred to an Immobilon membrane (Millipore; IPVH85R) After transfer, membranes were blocked in Tris-Buffered Saline (TBST) with w/ 1%BSA and then incubated overnight at 4oC in primary antibody. Antibodies used for western blot analysis are listed in Supplemental Table 2.

## RNA Sequencing

Human GBM cell lines (U118 and T98G) were treated with either vehicle control (DMSO) or 250nM NXP800 for 2 hours or 4 hours in biological triplicate (n=3). Cells were harvested using TrypLE (Gibco, Cat. #12605010), resuspended in DNA/RNA Shield (Zymo, Cat. #R1100), and submitted to Plasmidsaurus for total RNA isolation and bulk RNA sequencing (RNA-seq).

Quality control of raw single-end reads was evaluated using FastQC and MultiQC. Reads were aligned to the human reference genome (Ensembl GRCh38) using STAR, and gene-level counts were quantified using featureCounts (Subread package). Differential gene expression analysis was performed in R using DESeq2. Genes with an adjusted p-value < 0.05 and an absolute log2FC > 1.0 were identified as differentially expressed. Principal Component Analysis (PCA) and volcano plots were generated using ggplot2 and base R, respectively.

## Target Engagement Assay

U118 cells were cultured in 100 mm dishes in standard growth medium and treated with vehicle (DMSO) or 250nM of NXP800 or 5µM SISU-102 for 1 hour. Following treatment, the cells were rinsed with cold PBS and harvested in 350 µl of lysis buffer (40 mM HEPES, pH 7.5, 200mM NaCl, 0.4% NP-40) containing HALT™ Protease and Phosphatase Inhibitor Cocktail (Pierce, Cat. #1861281). Cell lysates were prepared by 10 seconds of sonication followed by centrifugation at 14,000 x g for 15 minutes. The protein concentration of the resulting supernatant was determined using a colorimetric assay (Protein Assay Dye; Bio-rad; 5000006) according to the manufacturer’s instructions. The resulting lysates were adjusted to 1mg/ml using complete lysis buffer, and 50 µl aliquots were subjected to a heat gradient: 25 °C for 2 minutes, then either 39.0°, 47.0°, 55.0°, 60.0°, 65.0°, 70.0°, 75.0°, 80.0°, and 90.0° for 3 minutes, followed by 2 minutes at 25 °C, and cooling to 4 °C. Following the temperature gradient, lysates were centrifuged at 20,000 × g for 20 minutes at 4 °C to remove insoluble material. Equal volumes of the supernatant were combined with BME (10%) + 4x Loading Dye (90%) and analyzed by western blot for HSF1 and GAPDH. Protein bands were quantified using FIJI (ImageJ) software and visualized using GraphPad Prism.

## Colony Formation Assay

Twenty-four hours after transfection, cells were seeded in triplicate into 6-well plates at 500- 1500 cells/well. After allowing cells to adhere to the plate (∼24hours), cells were then treated with indicated treatments. After 5-10 days, colonies were stained with 0.5% crystal violet.

Quantitative analysis of the colony formation assays was performed using FIJI (ImageJ) software to calculate colony number and area.

## Cell Viability Assay

Cells were seeded into a 96-well black bottom plate (Greiner; 655086) at 500-1000 cells/well. After incubation overnight, cells were treated with either DMSO or NXP800 for indicated lengths of time. CellTiter-Blue (CTB; Promega G8081) was added to each well at a 1:10 dilution at the end of treatment. Plates were then read using a Synergy H1 Microplate Reader at 560Ex/590Em. All measurements were performed in triplicate and normalized to the control.

Wells containing only medium served as the blank control.

## GCN2 Activity Signature

The GCN2 Activity Signature (GAS) was established using publicly available RNA-seq data (GSE145203). Differential expression analysis of approximately 40,000 genes initially identified 386 genes significantly upregulated by HFol (P>0.05; Log2FC>1). To ensure GCN2 specificity, genes that maintained HFol-responsiveness in the absence of GCN2 were excluded from the analysis. This filtering process yielded a refined 17-gene list designated as the GAS. HAS (30) and the AKT1 signature were previously established (34, 44). Z-scores were calculated from expression data (GSE196252, GSE188958, or PRJEB81176) to generate heatmaps in Morpheus (Broad Institute), and relative signature expression, quantified and analyzed using GraphPad Prism. Correlation between the HAS and GAS was further evaluated using The Cancer Genome Atlas (TCGA) patient data. Statistical significance was determined via one-way ANOVA followed by Tukey’s post-hoc test, with significance defined as P < 0.05.

## Statistical Tests

Statistical analysis was conducted using GraphPad Prism software (GraphPad Software, Inc., San Diego, CA), with all data expressed as the mean ± standard error of the mean (SEM) unless otherwise indicated. To determine statistical significance across experimental groups for the luciferase reporter, cell viability, and colony formation assay, analysis of variance (ANOVA) was employed followed by Tukey’s post-hoc test for multiple comparisons. Western blots presented in this study are representative of experiments performed in at least three independent biological replicates. For all quantitative assessments, including triplicate measurements of HSF1 activity and cell survival, a p-value<0.05 was considered statistical significance.

