## Supplementary figures and images for "NXP800 suppresses HSF1 through coordinated stress signaling and GCN2-dependent translational repression in glioblastoma"

### Supplemental Figures

Supplemental Figure 1

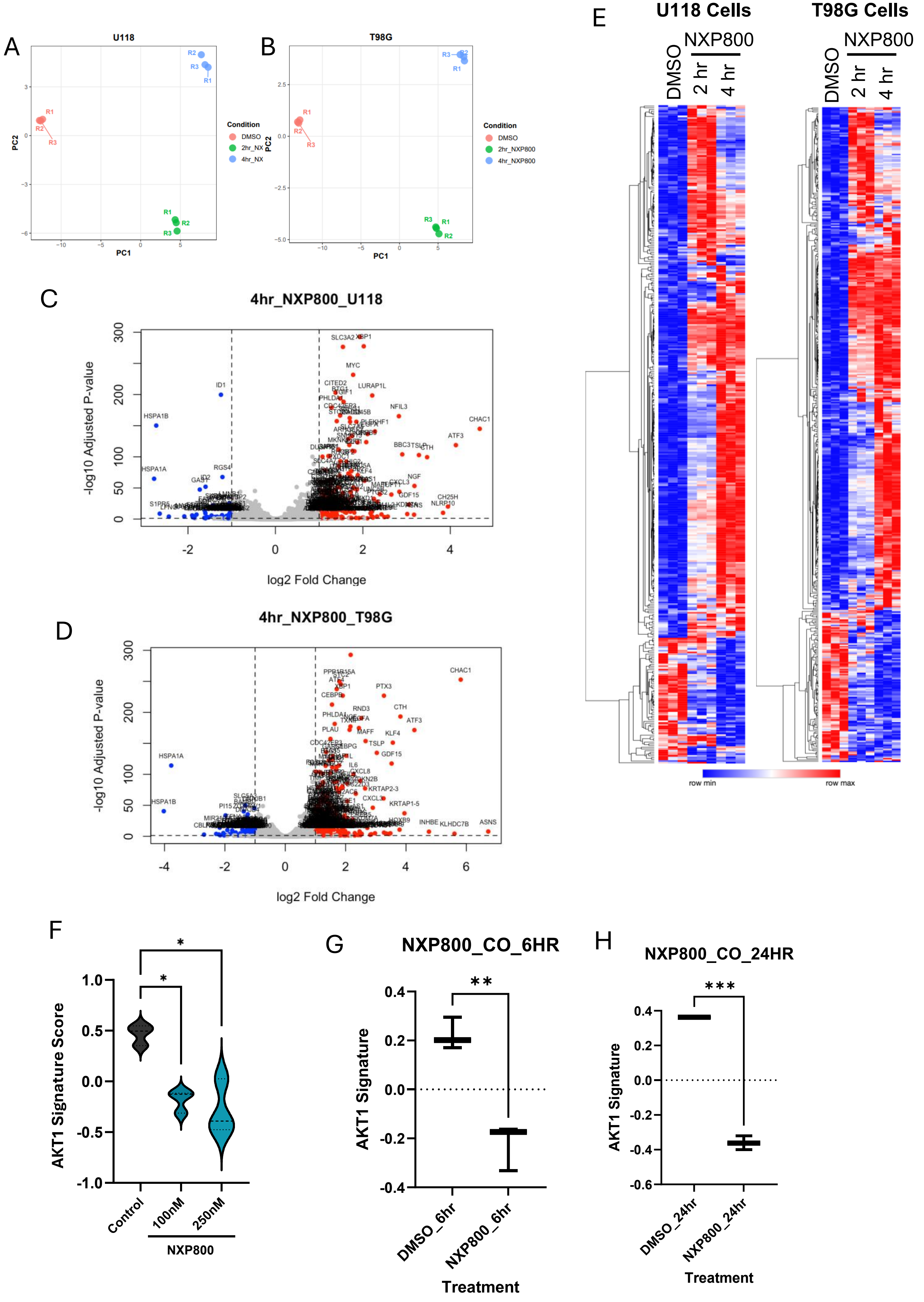

Supplemental Figure 2

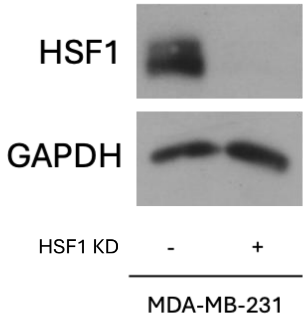

#### Supplemental Figure 3

A

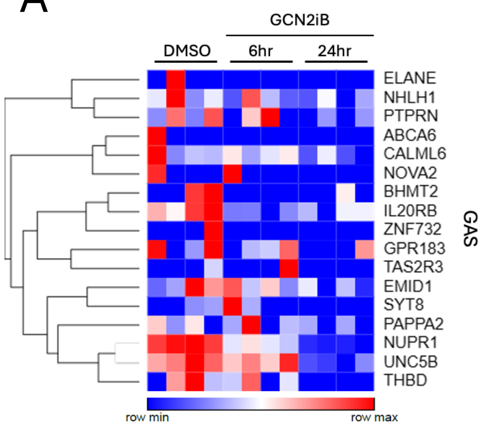


B

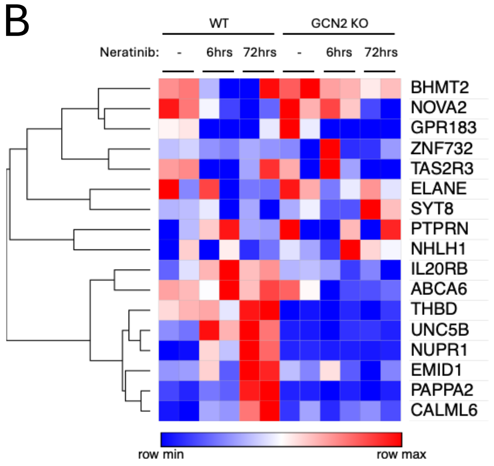


C

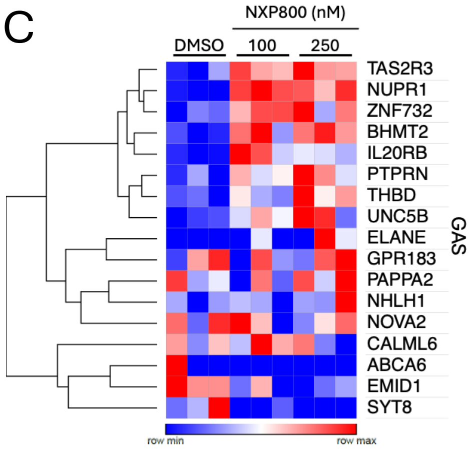
